# Split-QF system for fine-tuned transgene expression in *Drosophila*

**DOI:** 10.1101/451146

**Authors:** Olena Riabinina, Samuel W. Vernon, Barry J. Dickson, Richard A. Baines

**Affiliations:** Division of Neuroscience and Experimental Psychology, School of Biological Sciences, Faculty of Biology, Medicine and Health, Manchester Academic Health Science Centre, University of Manchester, Manchester, United Kingdom; Janelia Research Campus, HHMI, 19700 Helix Drive, Ashburn VA, 21407, USA

## Abstract

The Q-system is a binary expression system that works well across species. Here we report the development of a split-QF system that drives strong expression, is repressible by QS and inducible by quinic acid. The split-QF system is fully compatible with existing split-GAL4 and split-LexA lines for advanced intersectional experiments, thus greatly expanding the range of possible anatomical, physiological and behavioural assays in *Drosophila.*

## Main text

Binary expression systems GAL4/UAS^1^, LexA/LexAop^2^ and the Q-system^3-5^ allow labelling and functional manipulation of genetically defined subsets of cells in *Drosophila.* The split-GAL4 system^6-8^ allows expression of effectors to be limited to only a few cells by expressing a GAL4 DNA-binding domain (DBD) independently of a GAL4 activation domain (AD). A fully functional GAL4 is reconstituted only where the expression patterns of both subsets overlap. In practice, GAL4AD is often too weak and is replaced by p65AD or VP16AD to boost strength of expression^7,8^.

We reasoned that, since the QF2/QF2^w^ (a weaker version of QF2, with a mutated C-terminal^4^) transactivators of the Q-system are generally stronger than GAL4^4^, the split-QF system may function well in *Drosophila* by coupling QFDBD and QFAD^9^. This would allow the system to remain both repressible by QS and inducible by quinic acid (QA), in the same manner as the original Q-system. We have also previously developed chimeric GAL4QF and LexAQF transactivators^4^, which indicated that QFAD and QF2^w^AD are likely to function with GAL4DBD and LexADBD domains when brought together by leucine zippers.

To make the split-QF system compatible with existing split-GAL4 lines, we used the same leucine zippers^10^. We attached Zip-to QFDBD and Zip+ to QFAD and QF2^w^AD, defining the domains as previously reported^4^, and expressed these transgenes under control of the neuronal synaptobrevin promoter *nsyb* (**Fig 1a**). As expected, animals carrying *nsyb-QFDBD(attp40), nsyb-QFAD(attp2)* and *QUAS-mCD8-GFP* had strong GFP expression throughout their nervous system (**Fig 1B**). This expression was repressible by *tub-QS* and inducible by QA (**Supplementary Fig 1**). Similar, but weaker expression was observed with *nsyb-QFDBD* and *nsyb-QF2^w^AD* (**Fig 1B**). Both split transactivators appeared to have lower activity than the QF2 and QF2^w^ (**Fig 1B**).

**Figure 1.**
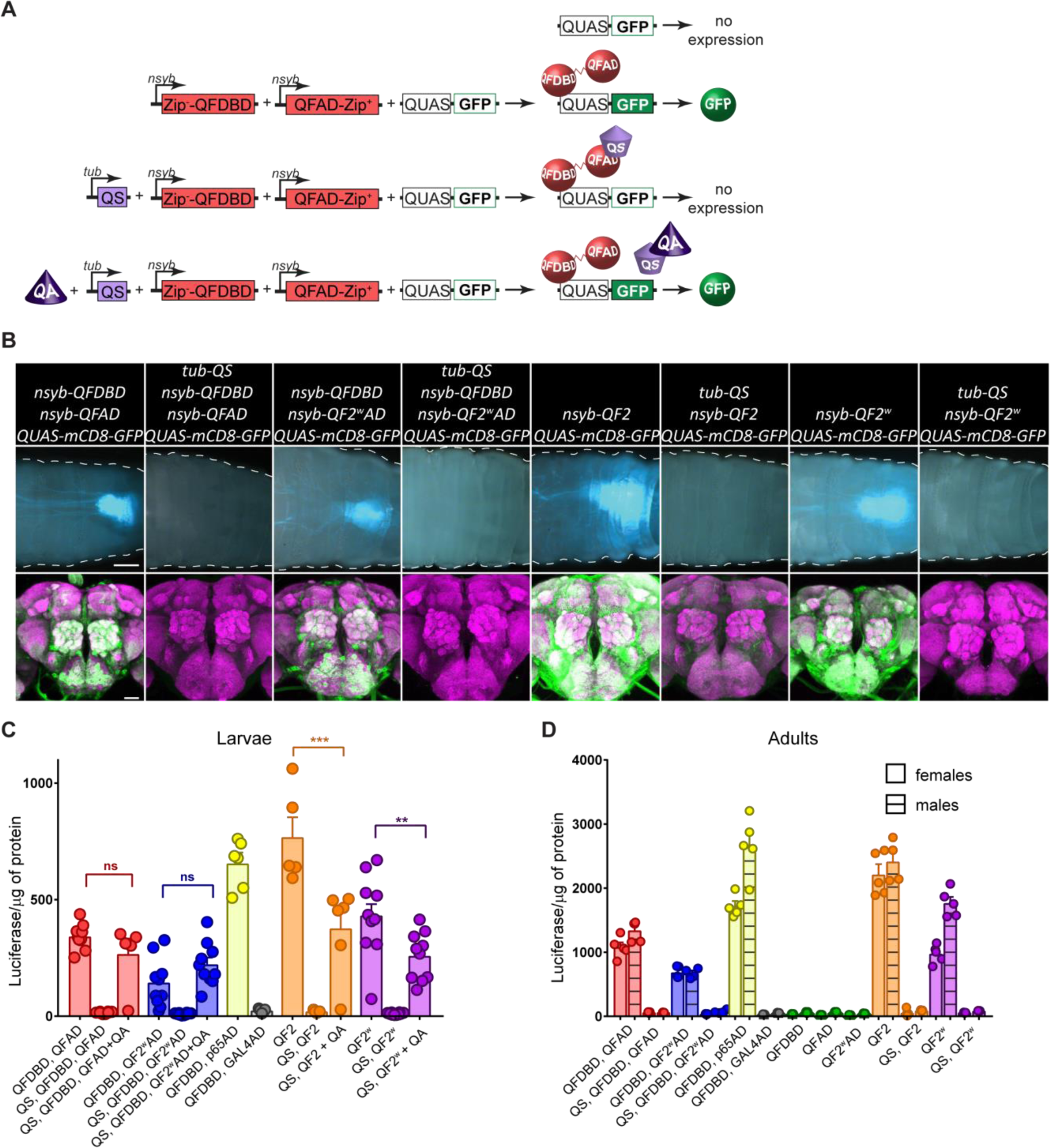
Quantification and validation of split-QF reagents. **A.** Schematics of the split-QF system. **B.** Pan-neuronal expression of GFP in larval (top; scale bar, 200μm) and adult (bottom; scale bar, 50μm) CNS by split-QF (first four columns) and Q-system (last four columns). **C, D.** Quantification of split-Q transactivators in larval (**C**) and adult (**D**) CNS by a luciferase assay. All split and full-length transactivators were driven by *nsyb*, while QS was driven by *tubulin.* Green data points show quantification for *nsyb-QFDBD, QUAS-luc; nsyb-QFAD, QUAS-luc* and *nsyb-QF2^w^AD, QUAS-luc* controls.

To compare QFAD and QF2^w^AD to existing p65AD and GAL4AD, we generated *nsyb-p65AD* (attp2) and *nsyb-GAL4AD* (attp2) flies, and expressed firefly luciferase pan-neuronally in larvae and adults (**Fig 1C,D**, **Supplementary Tables 1**,**2**). While relative expression levels varied between larvae (non-sexed) and male vs. female adults, QFAD was ∼2 times (p<0.01) stronger than QF2^w^AD, and ∼2 times (p<0.0001) weaker than p65AD. The GAL4AD was consistently weak. *tub-QS* provided strong repression of all original and split QF variants. We quantified the effect of QA de-repression in larvae only, because QA is effective only in sensory receptor neurons and the PI neurons in the adult brain^4^, presumably due to the glial blood-brain barrier^11^. QA feeding to *tub-QS, nsyb-QFDBD, nsyb-QFAD (QF2^w^AD)* larvae, that otherwise had very low expression, resulted in restoration of expression to the levels not significantly different from *nsyb-QFDBD, nsyb-QFAD (QF2^w^AD)* larvae (p=0.87 and p=0.62, respectively). These experiments demonstrate that the split-QF is fully functional, repressible and inducible, due to the strong activity of the QFAD and QF2^w^AD activation domains.

Next we asked whether QFAD and QF2^w^AD may be effectively used together with existing GAL4DBD lines, to provide an alternative to the currently used p65AD. Expression in larvae, driven by *elav-GAL4DBD* and *nsyb-QF2/QF2^w^AD*, is strong, QS-repressible and QA-inducible (**Fig 2A**, **top**). In adults the expression was strong and repressible in neurons consistent with the predicted expression pattern for each line, and QA-inducible in the olfactory and gustatory receptor neurons (**Fig 2A**, **bottom**, **Supplementary Fig 2**). To quantify the strength of expression, we used *elav-GAL4DBD* and the AD variants to drive luciferase in larval CNS. Note: the *elav-GAL4DBD, nsyb-p65AD* combination was lethal (**Fig 2B**, **Supplementary Table 3**). QFAD-induced expression was not significantly different from QF2^w^AD (p=0.16). In contrary to the experiments with split-QF (**Fig. 1C**), here QA resulted in restoration of expression to ∼20-35% of that of the un-repressed split transactivators (p<0.0001). To quantify expression levels in the adult CNS, we used *ChAT-GAL4DBD* to target cholinergic neurons and to avoid larval lethality, previously observed with *elav-GAL4DBD, nsyb-p65AD* (**Fig 2C**, **Supplementary Table 4**). QFAD-driven expression was comparable with QF2^w^AD (p>0.99) and ∼4 times weaker than p65AD (p<0.0001). *tub-QS* provided strong repression, not different from DBD-only or AD-only controls (p>0.99). These experiments demonstrate that QFAD and QF2^w^AD activation domains may be used together with GAL4DBD lines to provide a repressible and inducible, albeit weaker, alternative to p65AD.

**Figure 2.**
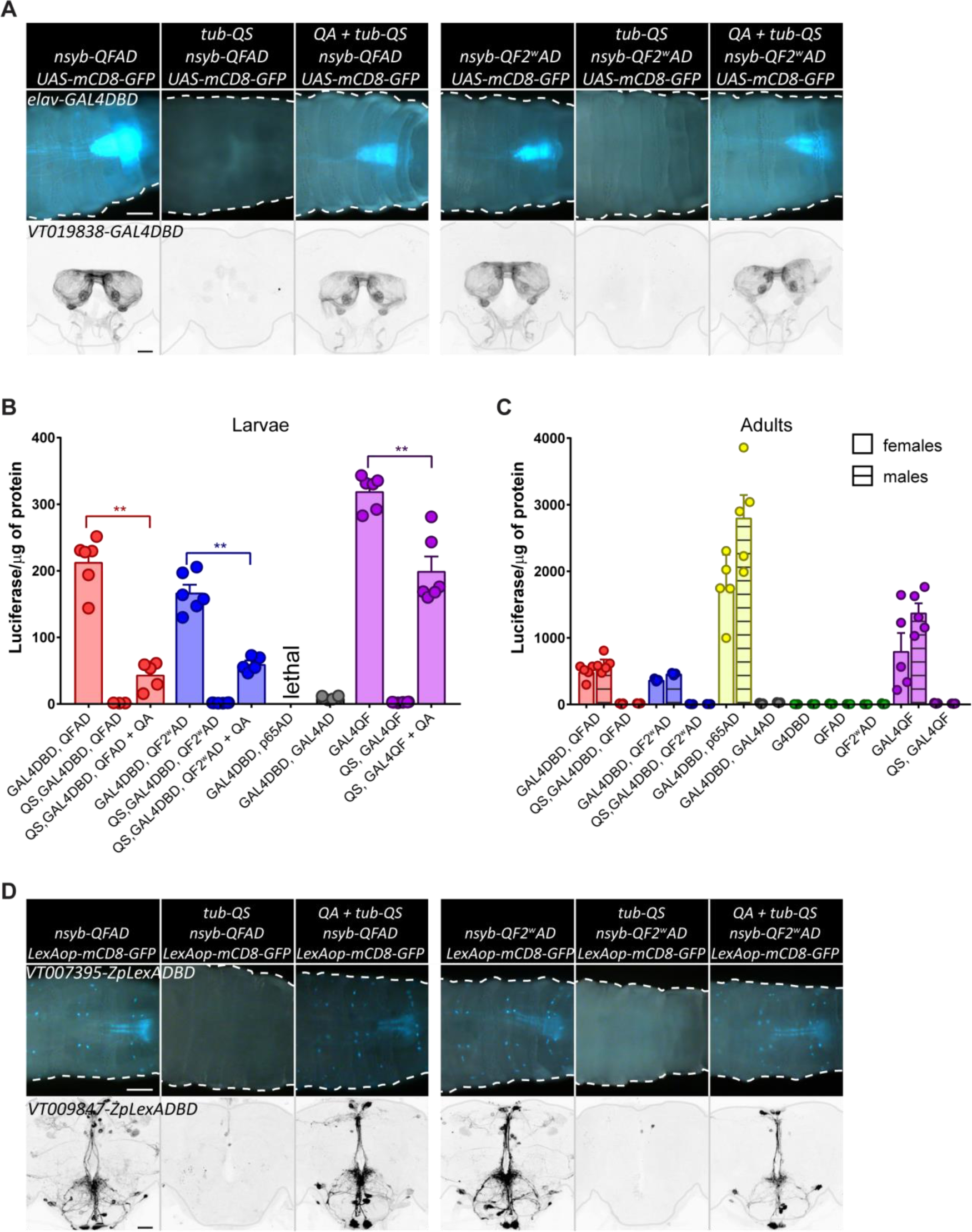
split-QF, split-GAL4 and split-LexA. A, top. Expression of GFP in larval CNS, driven by *elav-GAL4DBD* and *nsyb-QFAD* (3 left columns) or *nsyb-QF2^w^AD* (three right columns). Second and fifth columns show *tub-QS-induced* repression. Third and sixth columns show recovery of expression in larvae, grown on food with quinic acid. Scale bar, 200μm. A, bottom. Same as top, but driven by *VT019838-GAL4DBD* in adult CNS. Adults were fed with quinic acid for 5 days. Scale bar, 50μm. B. Quantification of relative strength of chimeric split transactivator in larval CNS. Genotypes were *elav-GAL4DBD, nsyb-QFAD* (red) or *elav-GAL4DBD, nsyb-QF2^w^AD* (blue) without (right) or with (middle) *tub-QS* and QA treatment (right). *elav-GAL4DBD, nsyb-GAL4AD* larvae (grey) had very low luciferase levels, while *elav-GAL4DBD, nsyb-p65AD* larvae did not survive. Purple bars show data from *nsyb-GAL4QF* larvae for comparison. C. Same as B, but in adult CNS. Males and females are quantified separately due to significantly different expression levels. Green data points show quantification for *elav-GAL4DBD, UAS-luc; nsyb-QFAD, UAS-luc* and *nsyb-QF2^w^AD, UAS-luc* controls. D, top. Expression of GFP in larval CNS, driven by *VT007395-LexADBD* and *nsyb-QFAD* (3 left columns) or *nsyb-QF2^w^AD* (three right columns). Second and fifth columns show tub-QS induced repression. Third and sixth columns show recovery of expression in the larvae, grown on food with quinic acid. Scale bar, 200μm. D, bottom. Same as top, but driven by *VT009847-LexADBD* in adult CNS. Adults were fed with quinic acid for 5 days. Scale bars, 50μm.

The QFAD and QF2^w^AD activation domains also work with split-LexA reagents in larval and adult CNS (**Fig 2D**, **Supplementary Figure 3**). Moreover, expression is both repressible and QA-inducible. Although we did not quantify strength of expression by luciferase (due to the unavailability of a LexAop-Luc reporter), it appears that QF2^w^AD domain works as well, or better, than QFAD in these experiments.

Next we asked how the QS repression compares with Killer-Zipper^12^ that silences split-GAL4 expression by driving GAL4DBD-Zip+ construct with the LexA/LexAop system (**Fig 3A**, **B**, **Supplementary Table 5**). We observed that QS-induced repression was stronger (p=0.0071 for *nsyb-QFDBD, nsyb-QFAD, KZip* vs *tub-QS, nsyb-QFDBD, nsyb-QFAD* females) or the same (all other genotypes, p>0.83) as a Killer-Zipper-induced equivalent. The use of QS for repression is thus more advantageous than Killer-Zipper because it requires fewer transgenes and does not recruit the LexA/LexAop system.

**Figure 3.**
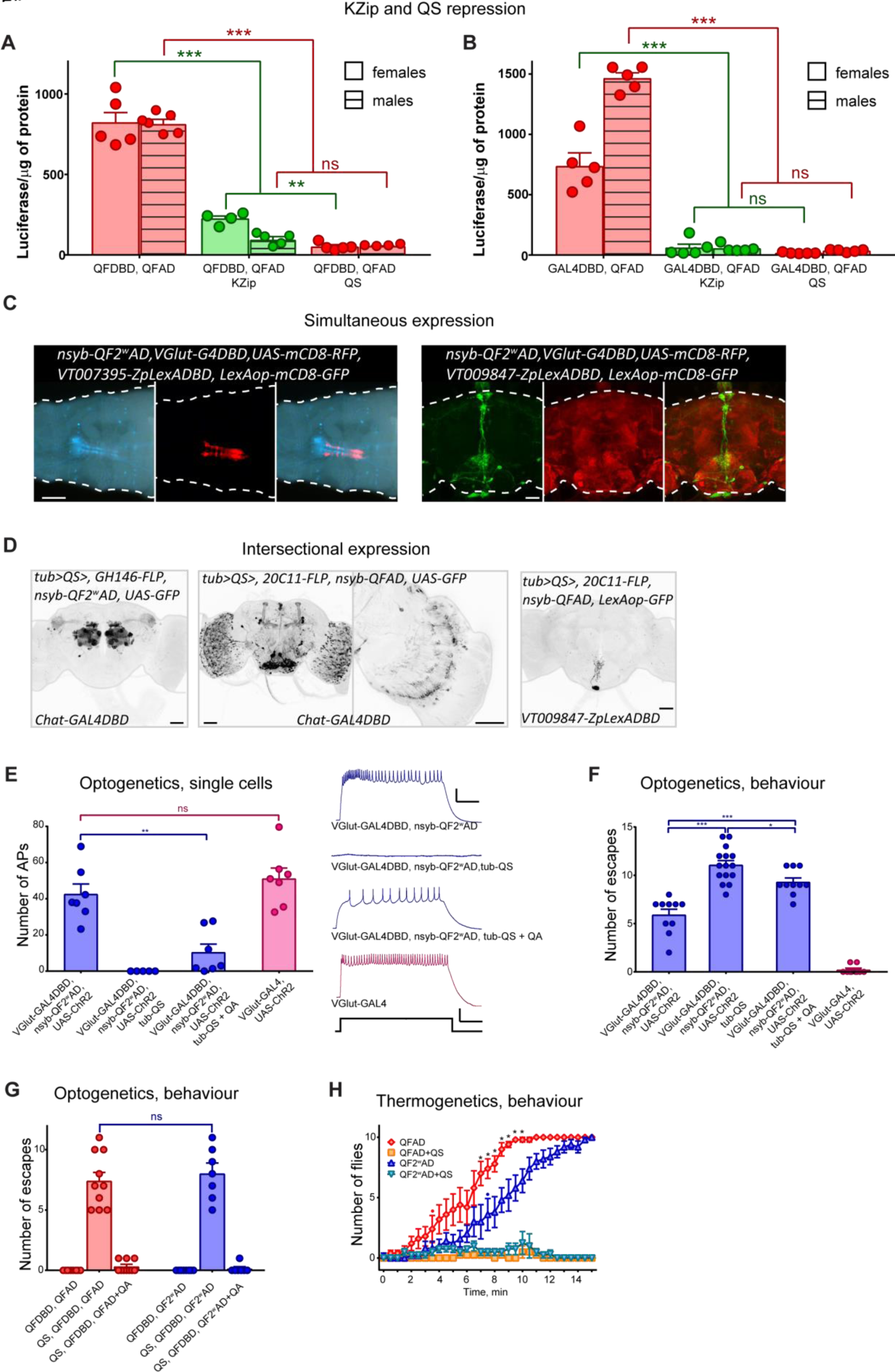
Applications of split-QF. A, B. Repression of expression by *Killer-Zipper*^12^ or *tub-QS.* Expression levels were quantified in adult flies using a luciferase assay. Genotypes of flies without repression were *nsyb-QFDBD, nsyb-QFAD, QUAS-Luc* (**A, right**) or *elav-GAL4DBD, nsyb-QFAD, UAS-Luc* (**B, right**). Killer-Zipper flies were *nsyb-QFDBD, nsyb-QFAD, nsyb-LexAQF, lexAop-KZip+, QUAS-Luc* **(A, middle, green)** or *elav-GAL4DBD, nsyb-QFAD, nsyb-LexAQF, lexAop-KZip+, UAS-Luc* **(B, middle, green).** QS flies were *tub-QS, nsyb-QFDBD, nsyb-QFAD, QUAS-Luc* (**A, left**) or *tub-QS, elav-GAL4DBD, nsyb-QFAD, UAS-Luc* (B, left). C. Simultaneous expression of RFP and GFP in independent neuronal subpopulations in larvae (left; scale bar, 200μm.) and adult (right; scale bar, 50μm) by *QF2^w^AD* forming functional transactivators with *GAL4DBD* and *LexADBD.* D. Intersectional expression, enabled by QS-repressible *GAL4DBD+QF/QF2^w^AD* and *LexADBD+QFAD* transactivators. GFP is expressed only in cells that 1) are expressing FLP or are progeny of cells that were expressing FLP; 2) are expressing G4DBD or LexADBD; 3) are expressing QFAD or QF2^w^AD. Third panel from the left shows a zoomed-in image of the z-stack of the brain, shown on the second panel. Scale bars, 50μm. **E**. Whole-cell patch-clamp recordings from aCC/PR2 motoneurons in third instar larvae of indicated genotypes, raised on food, supplemented with all-trans retinal. Depolarisation was elicited by blue light. Example traces are shown on the right. Scale Bars (Traces: 10mV/100ms, Stimulus: 2V/100ms). **F**. Escape assay of larvae with the same genotypes as in E. Each larva was given 2 mins to escape from a 113 mm^2^ area, lit by blue light (A.470 nm). Once the larva has completely left the lit area, it was returned into the area. G. Escape assay of *nsyb-QFDBD, nsyb-QFAD* larvae (red) and *nsyb-QFDBD, nsyb-QF2^w^AD* larvae (blue) with or without *tub-QS* and QA. H. Adult *nsyb-QFDBD, nsyb-QFAD, QUAS-shi^TS^* (red diamonds) and *nsyb-QFDBD, nsyb-QF2^w^AD, QUAS-shiTS* (dark blue upward triangles) flies were paralysed when placed in 33°C incubator at t=0 min. Flies that also had a *tub-QS* transgene (yellow squares and light-blue downward triangles) were not paralysed. The data shows the average number of flies (out of 10, ±SEM) at the bottom of the vial over time. Each graph is an average of n=5 repeats, apart from “QF2^w^AD+QS”, with n=4. Red dot and blue dot indicate the time point when the corresponding genotypes with and without QS became significantly different for the first time (t-test with Holm-Sidak correction for multiple comparisons). Stars indicate data points where *nsyb-QFDBD, nsyb-QFAD, QUAS-shi^TS^* and *nsyb-QFDBD, nsyb-QF2^w^AD, QUAS-shiTS* flies performed significantly differently (t-test with Holm-Sidak correction for multiple comparisons).

The split-QF system may be effectively used for simultaneous expression of UAS and LexAop transgenes: *QF2^w^AD*, when combined with *GAL4DBD* and *ZpLexADBD*, drives simultaneous expression of both *UAS-RFP* and *LexAop-GFP* (**Fig 3C**). To test the usability of split-QF for advanced intersectional experiments, we regulated expression of QS via the FLP-FRT system that, in turn, controlled the split-transactivators. As expected, intersection of *Chat-GAL4DBD, nsyb-QF2^w^AD* and *GH146-FLP* resulted in strong labelling of cholinergic olfactory projection neurons (**Fig 3D**, **left**). No labelling was observed when *Chat-GAL4DBD* was replaced by glutamatergic driver *VGlut-GAL4DBD* (not shown). Similarly, we observed expression throughout the brain and in the optic lobes in the cholinergic, but not glutamatergic (not shown), neurons that are targeted by *20C11-FLP*^13^ (**Fig 3D**, **middle**). Interestingly, intersection of *VT009847-ZpLexADBD, nsyb-QFAD* and *20C11-FLP* resulted in labelling only one SEZ neuron (**Fig 3D**, **right**). These experiments demonstrate that split-QF can effectively achieve simultaneous and intersectional expression, narrowing down expression patterns of split-GAL4, split-LexA and FLP lines.

We applied the split-QF system to study physiology and behaviour in *Drosophila.* We performed whole-cell patch-clamp recordings from aCC and RP2 motorneurons of third-instar larvae. Neuronal depolarisation was evoked through activation of UAS-ChR2^14^ expressed in all motoneurons by *VGlut-GAL4DBD, nsyb-QF2^w^AD* or, in controls, *VGlut-GAL4* (**Fig 3E**, **Supplementary Table 6**). The number of action potentials produced from *VGlut-GAL4DBD, nsyb-QF2^w^AD* larvae (42 ± 6 per 500ms) was not different from that in the GAL4 controls (51 ± 6, p=0.62). QS completely eliminated ChR2-induced depolarization in *tub-QS,VGlut-GAL4DBD, nsyb-QF2^w^AD* larvae (**Fig 3E**), while feeding larvae of the same genotype with QA partially restored depolarization and action potential count (10 ± 5), but significantly below the unrepressed levels of *VGlut-GAL4DBD, nsyb-QF2^w^AD* larvae (p=0.0016). These readouts of cellular activity are paralleled by behavioural phenotypes. We counted how many times (in 2 mins) larvae of these 4 genotypes escaped a blue light area (**Fig 3D**, **Supplementary Table 6**). As expected, larvae containing the QS transgene escaped most readily (11 ± 1.8 escapes), while feeding larvae with QA significantly reduced the number of escapes to 9.3 ± 1.3 (p=0.038), due to the seizure-like neuronal activity, elicited by ChR2 activation. *VGlut-GAL4DBD, nsyb-QF2^w^AD* were also able to escape (5.9 ± 0.6), but significantly less than the QS larvae (p<0.0001). *VGlut-GAL4* control larvae were unable to escape (0.2 ± 0.1).

We used the same assay to measure larval escape following activation of ChR2 driven pan-neuronally by split-QF (**Fig 3G**, **Supplementary Table 7**). Abolished mobility was observed in larvae that expressed ChR2 (0 ± 0 escapes in *nsyb-QFDBD, nsyb-QFAD* and *nsyb-QFDBD, nsyb-QF2^w^AD* larvaej, and in larvae that expressed QS and fed with QA (0.3 ± 0.2 and 0.1 ± 0.1 escapes for QFAD and QF2^w^AD, respectively). By contrast, QS-expressing larvae not fed with QA readily escaped the blue light area (7.4 ± 0.7 and 8.0 ± 0.8 escapes, respectively).

Finally, we assayed adult flies with pan-neuronal expression of *shibire^TS^* (**Fig 3H**, **Supplementary Table 8**). When placed in 33°C, *nsyb-QFDBD, nsyb-QFAD* flies became gradually paralysed as expected. The same effect was observed in *nsyb-QFDBD, nsyb-QF2^w^AD* flies, but took longer to develop, presumably due to the lower expression levels of *shibire^TS^.* When the expression of *shibire^TS^* was suppressed by *tub-QS*, no paralysis was observed.

These experiments demonstrate that split-QF may be used with or without split-GAL4 to direct expression of effectors in electrophysiological and behavioural assays.

In summary, we present a split-QF system that is applicable for advanced anatomical, behavioural and physiological manipulations in *Drosophila.* This system is fully compatible and complementary with the existing split-GAL4 and split-LexA lines and can greatly expand their use by making them QS-repressible and QA-inducible. In addition, combinations of split-QF with split-GAL4 and split-LexA systems can make extensive use of the available UAS and LexAop reporters.

## Acknowledgements

This work was supported by a Marie Curie Individual Fellowship (#701109) from the European Commission (OR). RAB was supported by funding from the Biotechnology and Biological Sciences Research Council (BB/L027690/1). Work on this project benefited from the Manchester Fly Facility, established through funds from the University and the Wellcome Trust (087742/Z/08/Z). Stocks obtained from the Bloomington Drosophila Stock Center (NIH P40OD018537) were used in this study.

## Author contributions

OR conceived of the project and designed most of the experiments. SWV and RB designed the electrophysiology experiments. OR and SWV performed the experiments. BJD provided unpublished reagents and suggestions. RB provided access to equipment and reagents. All authors contributed to drafting and revisions of the manuscript.

## Online methods

### Molecular biology

Plasmids were constructed by standard procedures including enzyme digestions, PCR and subcloning, using the In-Fusion HD Cloning System CE, Takara Bio Europe # 639636. Plasmid inserts were verified by DNA sequencing.

#### nsyb-nls::QFAD::Zip+ construct

1. 1) pattB-QF2-hsp70 plasmid (Addgene #46115) was digested with ZraI and EcoRI to remove Kozak-QF2 sequence.
2. 2) Kozak-nls sequence was PCR-amplified from pBPp65ADZpUw (Addgene #26234) with primers *ATC GAC AGC CGA ATT CAA CAT GGA TAA AGC GGA ATT A* (forward) and *ACG GTA TCG ATA GAC GTC CAA TTC GAC CTT TCT CTT C* (reverse).
3. 3) The PCR product was cloned into the digested vector by InFusion cloning.
4. 4) The cloning product was digested with ZraI
5. 5) QFAD sequence was PCR-amplified from pattB-QF2-hsp70 plasmid (Addgene #46115) with primers *AAG GTC GAA TTG GAC GTC CGT CAG TTG GAG CTA A* (forward) and *ACG GTA TCG ATA GAC AGA TCT CTG TTC GTA TGT ATT AAT GTC GGA GAA G* (reverse)
6. 6) The PCR product (5) was subcloned into (4) by InFusion cloning.
7. 7) (6) was digested with BglII
8. 8) The GGGGG-Zip+ sequence was PCR-amplified from pBPp65ADZpUw (Addgene #26234) with primers *ATA CGA ACA GAG ATC TGG AGG AGG TGG TGG AGG* (forward) and *ATC GAT AGA CAG ATC GGC CGG CCT TAC TTG CCG CCG CC* (reverse).
9. 9) The PCR product (8) was subcloned into the digested vector (7) by InFusion cloning.
10. 10) Product of (9) was digested with FseI and NotI to remove hsp70 terminator and to replace it with SV40 terminator
11. 11) SV40 terminator was PCR-amplfied from UAS-LUC-UAS-eYFP plasmid^15^ with primers *GGC AAG TAA GGC CGG CCG ATC TTT GTG AAG GAA CCT TAC* (forward) and *CCT CGA GCC GCG GCC GCG ATC CAG ACA TGA TAA GAT AC* (reverse).
12. 12) The PCR product (11) was subcloned into the vector (10) by InFusion cloning.

#### nsyb-nls::QF2^w^AD::Zip+ construct

1. 1) The *nsyb-nls::QFAD::Zip+* construct was digested with BglII and ZraI to remove QFAD.
2. 2) QF2^w^AD sequence was PCR amplified from from pattb-QF2-hsp70 (Addgene #46115) with primers AAG GTC GAA TTG GAC GTC CGT CAG TTG GAG CTC C (forward) and CAC CTC CTC CAG ATC TTT CTT CTT TTT GGT ATG TAT TAA TGT CGG AGA AGT TAC ATC C (reverse)
3. 3) The PCR product (2) was InFusino-cloned into (1).

#### nsyb-nls::p65AD::Zip+ construct

1. 1) The *nsyb-nls::QFAD::Zip+* construct was digested with FseI and ZraI to remove QFAD::Zip+ sequence.
2. 2) The p65AD::Zip+ sequence was PCR amplified from pBPp65ADZpUw (Addgene #26234) with primers *AAG GTC GAA TTG GAC GTC GGA TCC ACG CCG ATG* (forward) and *CTT CAC AAA GAT CGG CCG GCC TTA CTT GCC GCC GCC* (reverse).
3. 3) The PCR product (3) was InFusion-subcloned into (1).

#### nsyb-nls::GAL4AD::Zip+ construct

1. 1) The *nsyb-nls::QFAD::Zip+* construct was digested with BglII and ZraI to remove QFAD.
2. 2) The GAL4AD sequence was PCR amplified from pBPGAL4.2Uw-2 (Addgene #26227) with primers *AAG GTC GAA TTG GAC GTC GCC AAC TTC AAC CAG AGT GG* (forward) and *CAC CTC CTC CAG ATC TCT CCT TCT TTG GGT TCG GTG* (reverse).
3. 3) The PCR product (3) was InFusion-subcloned into (1).

#### nsyb-Zip-::QFDBD construct

1. 1) pattB-QF2-hsp70 plasmid (Addgene #46115) was digested with ZraI and EcoRI to remove Kozak-QF2 sequence.
2. 2) Kozak-Zip-GGGGGG sequence was PCR-amplified from pBPZpGAL4DBDUw (Addgene #26233) with primers *ATC GAC AGC CGA ATT CAA CAT GCT GGA GAT CCG C* (forward) and *ACG GTA TCG ATA GAC GTC ACC TCC ACC TCC ACC TCC* (reverse).
3. 3) The PCR product (3) was InFusion-subcloned into (1).
4. 4) (3) was digested with ZraI
5. 5) QFDBD was PCR-amplified from pattB-QF2-hsp70 plasmid (Addgene #46115) with primers *GGA GGT GGA GGT GAC GTC ATG CCA CCC AAG CG* (forward) and *ACG GTA TCG ATA GAC GGC CGG CCT TAG AGG AGG CGG GTA ATG C* (reverse).
6. 6) The PCR product (5) was InFusion-subcloned into (4).
7. 7) (6) was digested with FseI and NotI to remove hsp70 terminator and to replace it with SV40 terminator
8. 8) SV40 terminator was PCR-amplfied from UAS-LUC-UAS-eYFP plasmid^15^ with primers *CTC CTC TAA GGC CGG CCG ATC TTT GTG AAG GAA CCT TAC* (forward) and *CCT CGA GCC GCG GCC GCG ATC CAG ACA TGA TAA GAT AC* (reverse).
9. 9) The PCR product (8) was InFusion-subcloned into (7).

### Transgenic flies (new and existing)

New transgenic lines were generated by inserting *nsyb-QFDBD* construct in attp40 (II) and all *nsyb-AD* constructs into attp2 (III).

Other *Drosophila* stocks, used in this paper, were acquired from the Bloomington Stock Centre (indicated by # below) or were in personal stocks of the authors.

Figure 1: QUAS-mCD8-GFP (#30003), tub-QS (#52112), nsyb-QF2 (attp2, personal stocks, OR), nsyb-QF2^w^ (#51960), QUAS-Ppyr/Luc (#64773);

Figure 2: UAS-mCD8-GFP (personal stocks, OR), elav-GAL4DBD (derived from #23868), VT019838-GAL4DBD (#75177), ChAT-GAL4DBD (#60318), UAS-Luc (#64774), 13xLexAop2-mCD8-GFP (#32204), VT007395-ZpLexADBD (personal stocks, B.J.D.), VT009847-ZpLexADBD (personal stocks, B.J.D.);

Figure 3: nsyb-LexAQF (#51953), 13xLexAop2-KZip+ (#76253), VGlut-GAL4DBD (#60313), tub>QS> (#77125), GH146-FLP (gift of Christopher Potter, JHU), 20C11-FLP (#55766), UAS-ChR2 (gift of Stefan Pulver, St Andrews), VGlut-GAL4 (#60312), 10xQUAS-ChR2 (#52260), QUAS-shibire^TS^ (#30012).

Supplementary figures: R19F06-GAL4DBD (#69098), R53D01-GAL4DBD (#69075), VT059695-GAL4DBD (#73750), VT037031-ZpLexADBD (personal stocks, B.J.D.), VT043690-ZpLexADBD (personal stocks, B.J.D.).

### Immunohistochemistry and confocal imaging

Dissection and immunostaining of adult brains was done as described previously^4^. Briefly, on day 1 brains of 5-7 d.o. adult flies were dissected in ice-cold PBS, fixed at RT for 20 mins in 4% PFA in PBS+0.3% Triton (PBT), then washed in PBT at RT for 1.5-6h, blocked in 5% normal goat serum (NGS) in PBT for 30 mins and placed in primary antibody mix at 4°C for 3 nights on a shaker. On day 4, brains were washed in PBT at RT for 5-6h and placed in secondary antibody mix for 2 nights at 4°C on a shaker. On day 6, brains were washed in PBT for 5-6h and left overnight in approx. 50μl of Vectashield mounting solution without shaking. On day 7, brains were mounted in Vectashield on a microscope slide. The primary antibody mix contained rabbit anti-GFP (Invitrogen #A11122, 1:100), mouse nc82 (DSHB, 1:25) and 5% NGS in PBT. The secondary antibody mix contained Alexa Fluor 488 goat anti-rabbit (Invitrogen #A11034), Cy3 anti-mouse (Jackson Immunoresearch #115-165-062) and 5% NGS in PBT.

Images were acquired as z-stacks using a Leica SP8 upright confocal microscope equipped with HCX IRAPO L25x/0.95W water-immersion objective (Leica, Germany, 506323), at 512 × 512 pixel resolution with 1μm z steps. LAS X v3.5.2 software was used for image acquisition. Imaging settings (laser intensity, gain, etc.) were kept identical for groups of images that were compared to one another. Images were processed by taking maximum intensity projection, rotating and re-colouring in FIJI. Images shown are representative of 3-5 stainings for every genotype.

### Whole-animal imaging

Third-instar larvae were placed on a microscope slide and briefly put into a freezer to immobilize them. Images were taken on a Leica MZ10F zoom fluorescent scope equipped with a Leica DFC 420C camera, QImaging LED light source and LAS v.4 software. The white balance was adjusted automatically by taking an image of a white sheet of paper before experimental images. Identical settings were used to take images that are compared to each other. Images shown are representative of 3-5 for every genotype.

### Quinic acid feeding

For larval experiments, gravid females were allowed to lay eggs in vials containing standard fly medium, supplemented with QA, and larvae remained in the vials until they reached wall-climbing 3^rd^ instar stage. For adult experiments, flies were raised on standard fly medium and were transferred into vials with QA at 2-3 d.o., for 5 days, at which point they were dissected. To make QA stock, 8 g of QA (Sigma #138622) was dissolved in 40 ml ddH_2_0 and adjusted to pH7 with 5M NaOH, bringing the total stock volume to 50 ml. 1.6ml/vial of this solution was thoroughly mixed into standard fly medium for larval or adult experiments.

### Luciferase assay

Each experiment assayed 9-30 larvae or 9-15 adult flies per genotype, in groups of three. 3^rd^ instar larvae or 1-2d.o. adult flies were placed in a 1.5 ml Eppendorf tube and stored in-80°C until all samples for a given experiment were collected. A Dual-Luciferase Reporter Assay system (Promega, E1910) was used for the experiments. Samples were homogenised in 200μl of Passive lysis buffer (Promega, E194A) per tube, and kept on ice for at least 10 mins. Then the tubes were centrifuged for 5 mins at 13.4k rpm, and supernatant transferred to new tubes. 30μl of supernatant from each tube were mixed with 30μl of Luciferase assay substrate (Promega, E151A), reconstituted in Luciferase assay buffer (Promega, E195A), per well of a 96-well plate and luminescence was measured immediately on a TECAN GENios plate reader, running XFluor 4 macros for Excel. We used 300ms exposure for adult samples and 600ms exposure for larval samples. We collected 3-10 measurements per experiment per genotype. The luciferase luminescence values were normalised by the amount of protein contained in the samples, to account for possible differences in the size of larvae and adults. For protein assay, 1.5ul of supernatant was mixed with 100ul of Protein assay reagent (BioRAD, #500-0006) and light absorbance measured after 20 mins on a FLUOstar Omega platereader (BMG LABTECH), running Omega software v. 1.3. Two independent samples were measured per each supernatant tube. The absorbance values were converted into mg/ml of protein by measuring a calibration curve with BSA dilutions (NEB, #B90015). Each relative luminescence (RL) data point, presented on the graphs (**Fig 1C-D, 2B-C, 3A-B**) was calculated as follows:

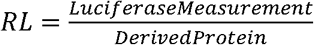, where 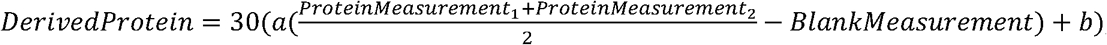, *a* and *b* parameters were obtained from the best linear fit to the calibration curve, plotted as (*average of 3 calibration measurement for a given dilution of BSA)-blank measurement* vs *dilution of BSA in mg/ml.* 4-6 independent RL values were collected per genotype in each experiment. The genotypes are presented in Figures as mean±SEM and were compared with 1-way ANOVA (larvae) or 2-way ANOVA (adults) with Sidak’s multiple comparisons test.

We have observed significant differences between measurement of adult males and females for some genotypes, arising from a consistently higher amount of protein per adult female. These differences were never observed for male and female larvae (data not shown). Thus, we present adult data separately for males and females.

### Larval whole-cell patch-clamp recordings

Larvae were grown in the dark on standard fly medium, supplemented with 100μl/vial of 0.1M all trans-retinal (Sigma, #R2500) in 100% EtOH. Recordings were performed at room temperature (20-22°C). Third-instar larvae were dissected in external saline (in mM: 135 NaCl, 5 KCl, 4 MgCl2-6H2O, 2 CaCl2-2H2O, 5 N-Tris[hydroxymethyl]methyl-2-aminoethanesulfonic acid, and 36 sucrose, pH 7.15). The CNS was removed and secured to a Sylgard (Dow-Corning, Midland, Michigan, USA)-coated cover slip using tissue glue (GLUture; WPI, Hitchin, UK). The glia surrounding the CNS was partially removed using protease (1% type XIV; Sigma, Dorset, UK) contained in a wide-bore (15 μm) patch pipette. Whole cell recordings were carried out using borosilicate glass electrodes (GC100TF-10; Harvard Apparatus, Edenbridge, UK), fire-polished to resistances of between 10-14 MΩ. The aCC/RP2 motoneurons were identified by soma position within the ventral nerve cord. When needed, cell identity was confirmed after recording by filling with 0.1% Alexa Fluor 488 hydrazyde sodium salt (Invitrogen, Carlsbad, California, USA), included in the internal patch saline (in mM: 140 potassium gluconate, 2 MgCl2-6H2O, 2 EGTA, 5 KCl, and 20 HEPES, pH 7.4). Mecamylamine (1 mM, M9020, Sigma, Dorset, UK) was included in the external saline to block endogenous excitatory cholinergic mediated currents to aCC/RP2 motoneurons and neuronal depolarisation was elicited through UAS-ChR2^14^ (A470 nm, 500ms, light intensity 9.65 mW/cm^2^ before reaching the LUMPlanFI 60×/0.9W Olympus objective) expressed in all motoneurons by VGlut promoter. Recordings were made using a MultiClamp 700B amplifier. Cells were held at-55 mV and recordings were sampled at 20 kHz and lowpass filtered at 10 kHz, using pClamp 10.6 (Molecular Devices, Sunnyvale, CA). Only neurons with an input resistance of > 500 MO were accepted for analysis. 8 recordings were taken per cell, average action potential number per 500ms light pulse calculated. Data in **Fig 3E** are presented as mean±SEM, and were compared with a 1-way ANOVA with Sidak’s multiple comparisons test.

### Larval escape assays

Individual 3^rd^ instar larvae were assayed at RT (20-22C) in a 9cm petri dish that contained a thin layer of 1% agarose to prevent desiccation. The petri dish was placed under the Leica MZ16F zoom fluorescent microscope with Plan 1.0x lens, fluorescent light source and a GFP filter cube (λ470 nm). Light intensity measured 9.87 mW/cm^2^ when completely zoomed out. Zoom 5 was used for experiments. Larvae were filmed using a uEye UI-233xSE-C camera with uEye Cockpit software, and data was stored in *.avi format. Each larvae was allowed to crawl in the Petri dish for 2 mins, before it was placed for 2 mins into a 113mm^2^ area, illuminated by blue light. Wild-type larvae naturally avoid bright blue light and crawl away, however, larvae with ChR2 expressed in motoneurons (**Fig 3F**) or pan-neuronally (**Fig 3G**) are impaired in their ability to escape. A larva was returned into the blue light area immediately after the larva had completely left the illuminated area. We counted the number of escapes during a 2 min period. 7-15 larvae were assayed per genotype. The data is shown as mean±SEM. The genotypes were compared with 1-way ANOVA with Sidak’s multiple comparison test.

### Adult behavioural assay

Adult male and female 5-7d.o. flies were assayed in groups of 10 (N=4−5 groups per genotype) in clean empty standard fly vials. Flies were placed in a cooled incubator, set to 33°C, and videorecorded at 5 fps using a uEye camera UI-233xSE-C, controlled by uEye Cockpit software. The data was stored in *.avi format. The number of flies on the bottom of each vial was manually counted at 30s intervals. The data is shown as mean±SEM, and was analysed with multiple t-tests with Holm-Sidak correction.

**Supplemental Figure 1.**
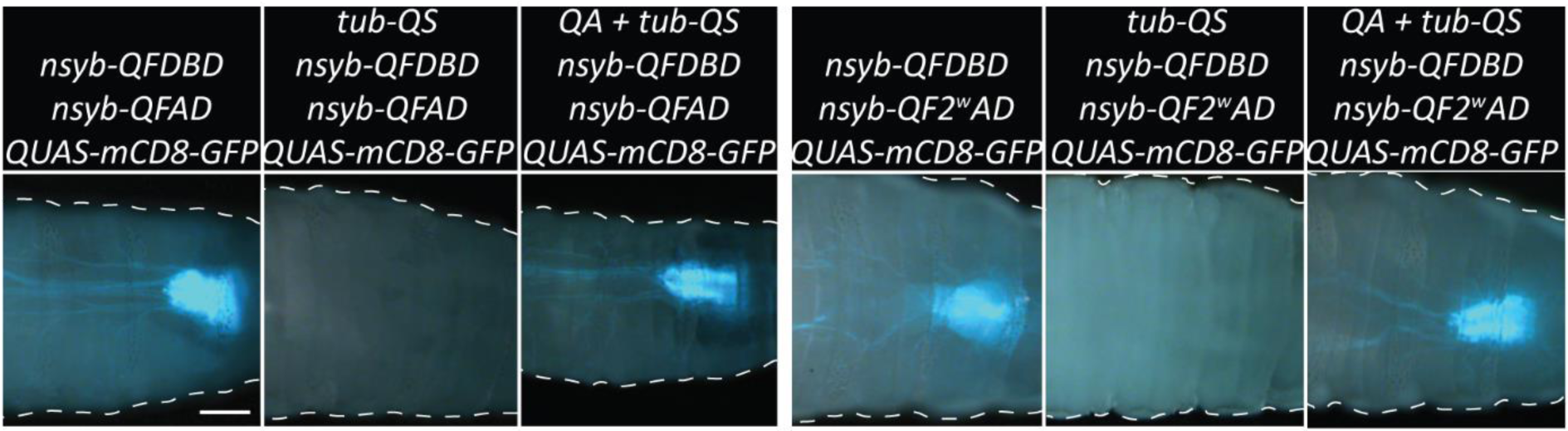
Quantification and validation of split-QF reagents. Pan-neuronal expression of GFP in the larval CNS by split-QF. Panels 1,2, 4 and 5 are the same as in Fig 1. Panels 3 and 6 show QA-induced de-repression. Scale bars, 200μm.

**Supplemental Figure 2.**
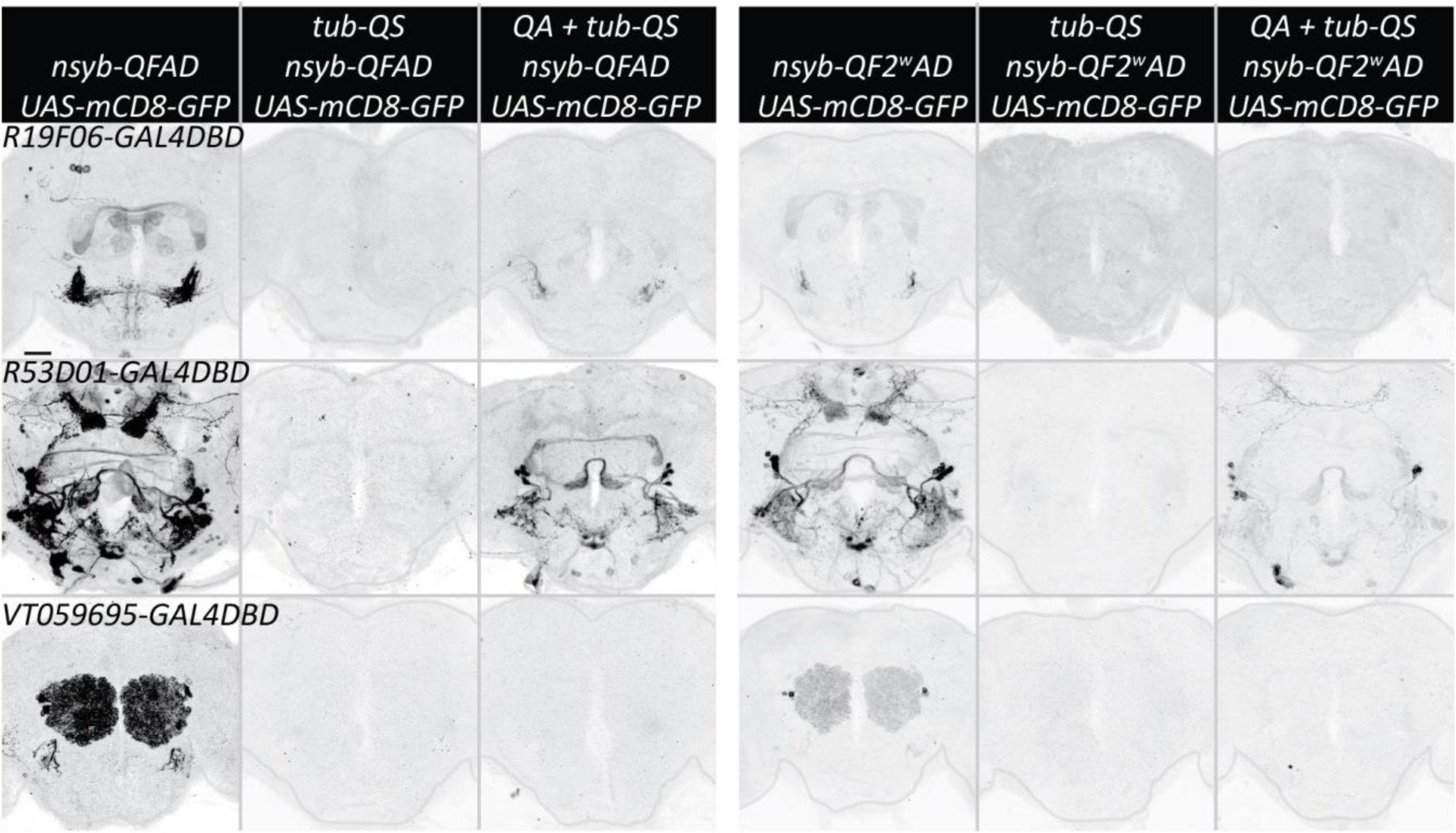
split-QF and split-GAL4. Expression of GFP in adult CNS, driven by *R19F06-GAL4DBD* (top), *R53D01-GAL4DBD* (middle), *VT059695-GAL4DBD* (bottom) and *nsyb-QFAD* (3 left columns) or *nsyb-QF2^w^AD* (three right columns). Second and fifth columns show tub-QS-induced repression. Third and sixth columns show recovery of expression in adults that were fed quinic acid for 5 days. Scale bar, 50μm.

**Supplemental Figure 3.**
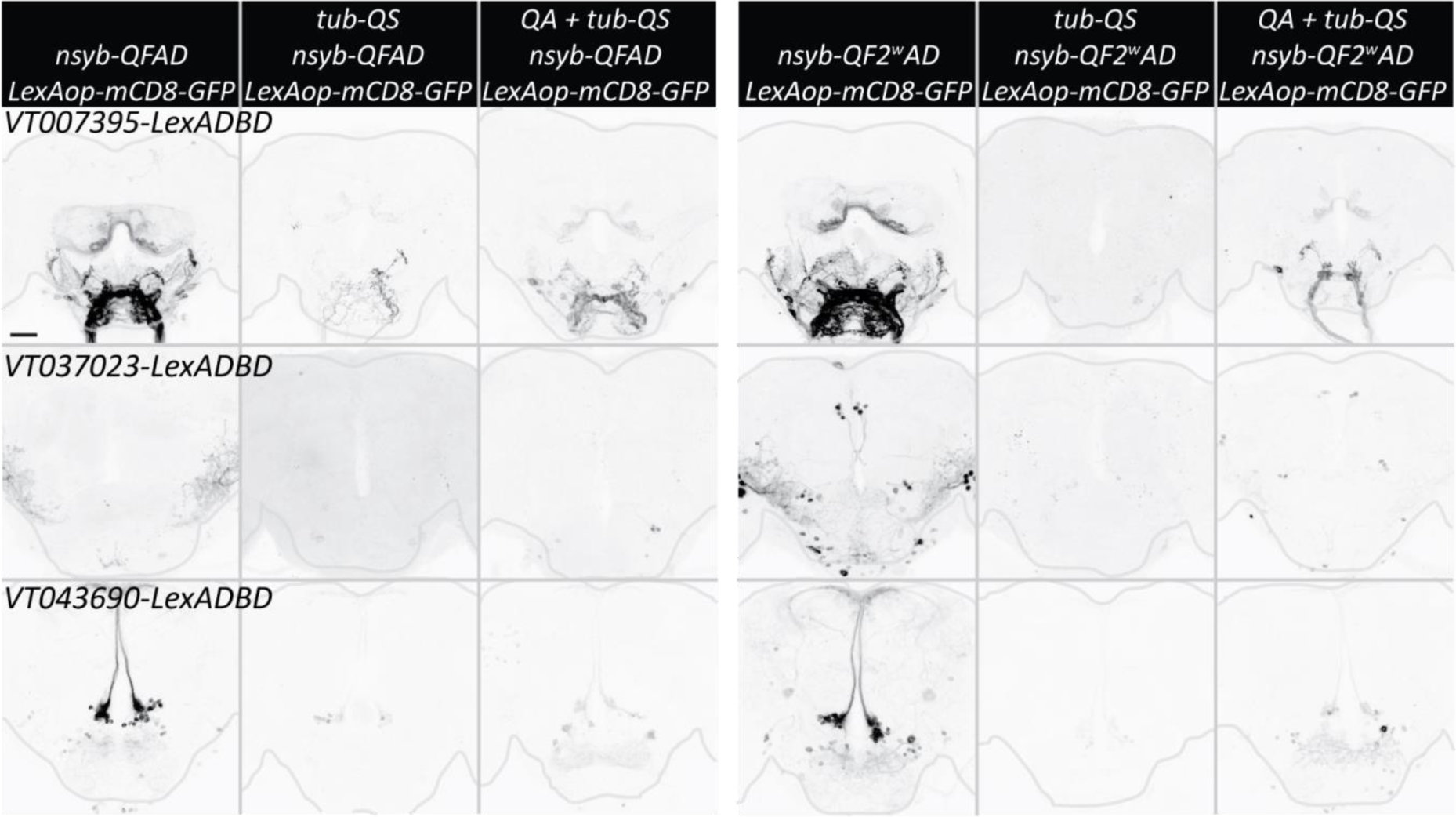
split-QF and split-LexA. Expression of GFP in adult CNS, driven by *VT007395-LexADBD* (top), *VT037023-LexADBD* (middle), *VT043690-LexADBD* (bottom) and *nsyb-QFAD* (3 left columns) or *nsyb-QF2^w^AD* (three right columns). Second and fifth columns show *tub-QS-* induced repression. Third and sixth columns show recovery of expression in adults that were fed quinic acid for 5 days. Scale bar, 50μm.

**Supplementary Table 1.**
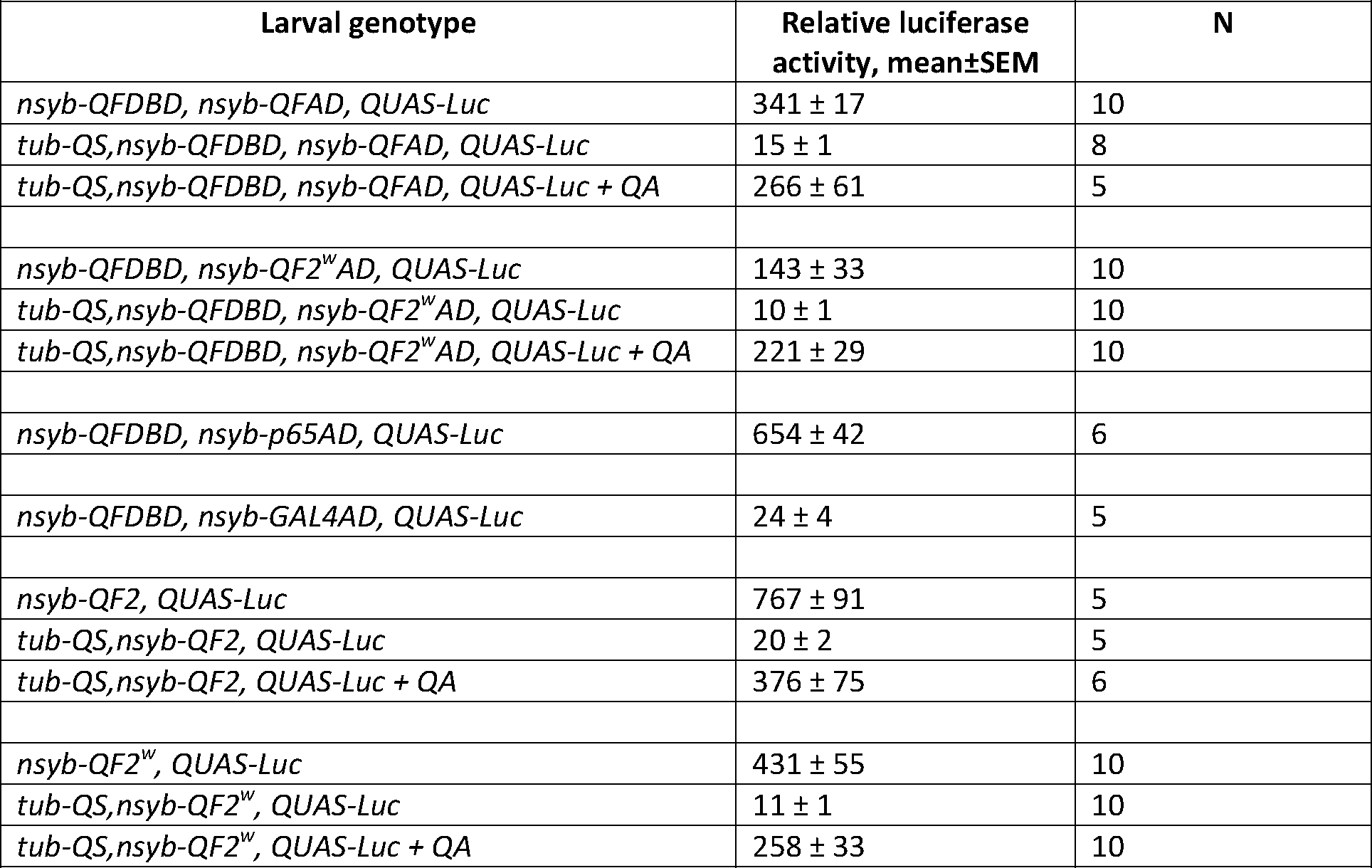
Quantification of expression strength of split-QF reagents in larvae

| Larval genotype | Relative luciferase activity, mean±SEM | N |
| --- | --- | --- |
| *nsyb-QFDBD, nsyb-QFAD, QUAS-Luc* | 341 ± 17 | 10 |
| *tub-QS,nsyb-QFDBD, nsyb-QFAD, QUAS-Luc* | 15 ± 1 | 8 |
| *tub-QS,nsyb-QFDBD, nsyb-QFAD, QUAS-Luc + QA* | 266 ± 61 | 5 |
| *nsyb-QFDBD, nsyb-QF2<sup>w</sup>AD, QUAS-Luc* | 143 ± 33 | 10 |
| *tub-QS,nsyb-QFDBD, nsyb-QF2<sup>w</sup>AD, QUAS-Luc* | 10 ± 1 | 10 |
| *tub-QS,nsyb-QFDBD, nsyb-QF2<sup>w</sup>AD, QUAS-Luc + QA* | 221 ± 29 | 10 |
| *nsyb-QFDBD, nsyb-p65AD, QUAS-Luc* | 654 ± 42 | 6 |
| *nsyb-QFDBD, nsyb-GAL4AD, QUAS-Luc* | 24 ± 4 | 5 |
| *nsyb-QF2, QUAS-Luc* | 767 ± 91 | 5 |
| *tub-QS,nsyb-QF2, QUAS-Luc* | 20 ± 2 | 5 |
| *tub-QS,nsyb-QF2, QUAS-Luc + QA* | 376 ± 75 | 6 |
| *nsyb-QF2<sup>w</sup>, QUAS-Luc* | 431 ± 55 | 10 |
| *tub-QS,nsyb-QF2<sup>w</sup>, QUAS-Luc* | 11 ± 1 | 10 |
| *tub-QS,nsyb-QF2<sup>w</sup>, QUAS-Luc + QA* | 258 ± 33 | 10 |

**Supplementary Table 2.**
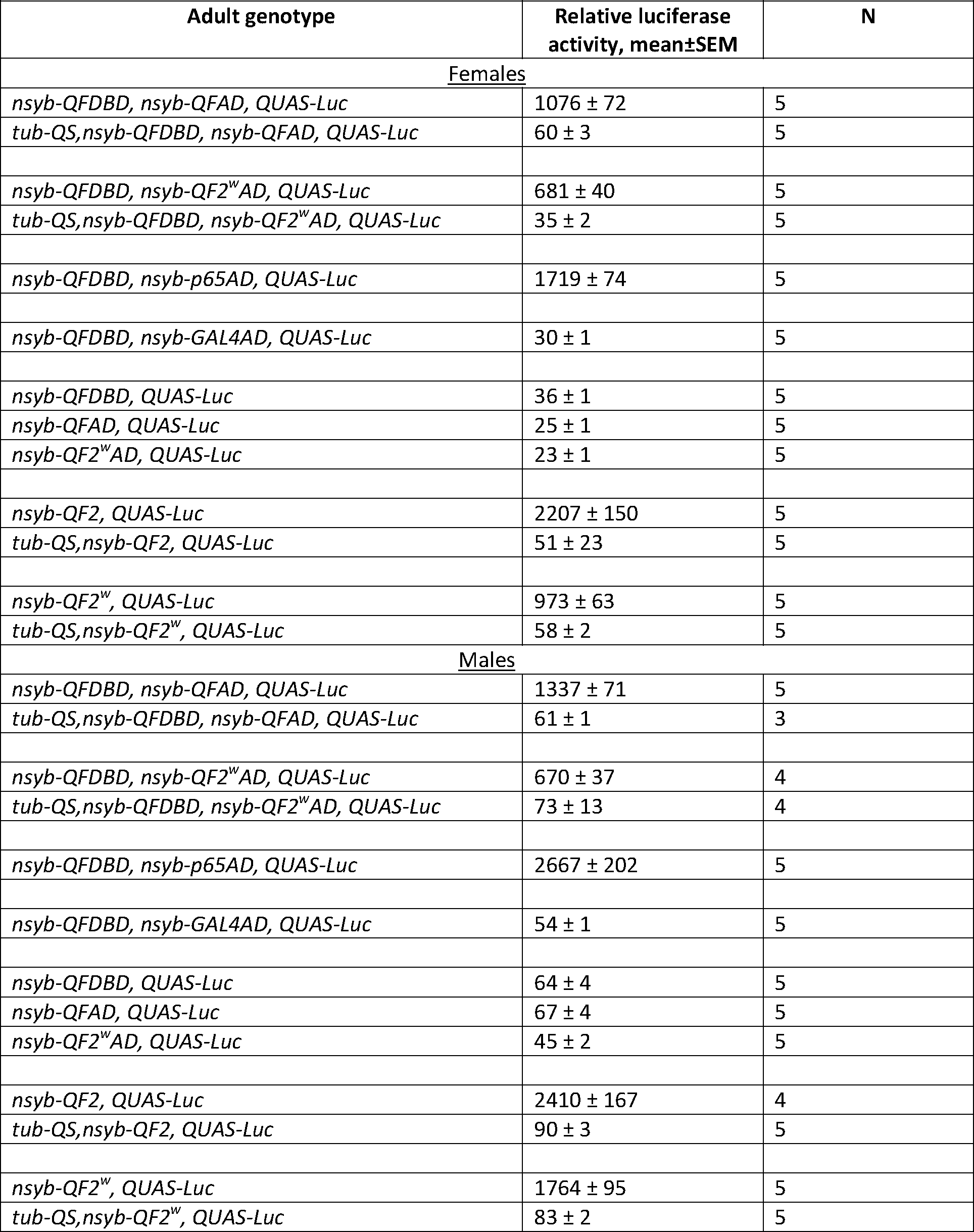
Quantification of expression strength of split-QF reagents in adults

| Adult genotype | Relative luciferase activity, mean±SEM | N |
| --- | --- | --- |
| <u>Females</u> |  |  |
| *nsyb-QFDBD, nsyb-QFAD, QUAS-Luc* | 1076 ± 72 | 5 |
| *tub-QS,nsyb-QFDBD, nsyb-QFAD, QUAS-Luc* | 60 ± 3 | 5 |
| *nsyb-QFDBD, nsyb-QF2<sup>W</sup>AD, QUAS-Luc* | 681 ± 40 | 5 |
| *tub-QS,nsyb-QFDBD, nsyb-QF2<sup>W</sup>AD, QUAS-Luc* | 35 ± 2 | 5 |
| *nsyb-QFDBD, nsyb-p65AD, QUAS-Luc* | 1719 ± 74 | 5 |
| *nsyb-QFDBD, nsyb-GAL4AD, QUAS-Luc* | 30 ± 1 | 5 |
| *nsyb-QFDBD, QUAS-Luc* | 36 ± 1 | 5 |
| *nsyb-QFAD, QUAS-Luc* | 25 ± 1 | 5 |
| *nsyb-QF2<sup>W</sup>AD, QUAS-Luc* | 23 ± 1 | 5 |
| *nsyb-QF2, QUAS-Luc* | 2207 ± 150 | 5 |
| *tub-QS,nsyb-QF2, QUAS-Luc* | 51 ± 23 | 5 |
| *nsyb-QF2<sup>W</sup>, QUAS-Luc* | 973 ± 63 | 5 |
| *tub-QS,nsyb-QF2<sup>W</sup>, QUAS-Luc* | 58 ± 2 | 5 |
| <u>Males</u> |  |  |
| *nsyb-QFDBD, nsyb-QFAD, QUAS-Luc* | 1337 ± 71 | 5 |
| *tub-QS,nsyb-QFDBD, nsyb-QFAD, QUAS-Luc* | 61 ± 1 | 3 |
| *nsyb-QFDBD, nsyb-QF2<sup>W</sup>AD, QUAS-Luc* | 670 ± 37 | 4 |
| *tub-QS,nsyb-QFDBD, nsyb-QF2<sup>W</sup>AD, QUAS-Luc* | 73 ± 13 | 4 |
| *nsyb-QFDBD, nsyb-p65AD, QUAS-Luc* | 2667 ± 202 | 5 |
| *nsyb-QFDBD, nsyb-GAL4AD, QUAS-Luc* | 54 ± 1 | 5 |
| *nsyb-QFDBD, QUAS-Luc* | 64 ± 4 | 5 |
| *nsyb-QFAD, QUAS-Luc* | 67 ± 4 | 5 |
| *nsyb-QF2<sup>W</sup>AD, QUAS-Luc* | 45 ± 2 | 5 |
| *nsyb-QF2, QUAS-Luc* | 2410 ± 167 | 4 |
| *tub-QS,nsyb-QF2, QUAS-Luc* | 90 ± 3 | 5 |
| *nsyb-QF2<sup>W</sup>, QUAS-Luc* | 1764 ± 95 | 5 |
| *tub-QS,nsyb-QF2<sup>W</sup>, QUAS-Luc* | 83 ± 2 | 5 |

**Supplementary Table 3.** Quantification of expression strength of split-QF + split-GAL4 reagents in larvae

| Larval genotype | Relative luciferase activity, mean±SEM | N |
| --- | --- | --- |
| *elav-GAL4DBD, nsyb-QFAD, UAS-Luc* | 213 ± 16 | 6 |
| *tub-QS, elav-GAL4DBD, nsyb-QFAD, UAS-Luc* | 1.4 ± 0.1 | 3 |
| *tub-QS, elav-GAL4DBD, nsyb-QFAD, UAS-Luc + QA* | 44 ± 9 | 5 |
| *elav-GAL4DBD, nsyb-QF2<sup>w</sup>AD, UAS-Luc* | 167 ± 12 | 6 |
| *tub-QS, elav-GAL4DBD, nsyb-QF2<sup>w</sup>AD, UAS-Luc* | 1.6 ± 0.1 | 6 |
| *tub-QS, elav-GAL4DBD, nsyb-QF2<sup>w</sup>AD, UAS-Luc + QA* | 60 ± 5 | 5 |
| *elav-GAL4DBD, nsyb-p65AD, UAS-Luc* | lethal | 0 |
| *elav-GAL4DBD, nsyb-GAL4AD, UAS-Luc* | 9 ± 1 | 6 |
| *nsyb-GAL4QF, UAS-Luc* | 319 ± 10 | 6 |
| *tub-QS,nsyb-GAL4QF, UAS-Luc* | 2.5 ± 0.2 | 6 |
| *tub-QS,nsyb- GAL4QF, UAS-Luc + QA* | 199 ± 21 | 6 |

**Supplementary Table 4.** Quantification of expression strength of split-QF + split-GAL4 reagents in adults

| Adult genotype | Relative luciferase activity, mean±SEM | N |
| --- | --- | --- |
| <u>Females</u> |  |  |
| *ChAT-GAL4DBD, nsyb-QFAD, UAS-Luc* | 488 ± 50 | 5 |
| *tub-QS, ChAT-GAL4DBD, nsyb-QFAD, UAS-Luc* | 10 ± 3 | 5 |
| *ChAT-GAL4DBD, nsyb-QF2<sup>w</sup>AD, UAS-Luc* | 366 ± 8 | 5 |
| *tub-QS, ChAT-GAL4DBD, nsyb-QF2<sup>w</sup>AD, UAS-Luc* | 6 ± 1 | 5 |
| *ChAT-GAL4DBD, nsyb-p65AD, UAS-Luc* | 1763 ± 217 | 5 |
| *ChAT-GAL4DBD, nsyb-GAL4AD, UAS-Luc* | 12 ± 4 | 5 |
| *ChAT-GAL4DBD, UAS-Luc* | 4.2 ± 0.6 | 5 |
| *nsyb-QFAD, UAS-Luc* | 8 ± 2 | 5 |
| *nsyb-QF2<sup>w</sup>AD, UAS-Luc* | 2.8 ± 0.4 | 5 |
| *nsyb-GAL4QF, UAS-Luc* | 798 ± 274 | 5 |
| *tub-QS, nsyb-GAL4QF, UAS-Luc* | 16 ± 2 | 5 |
| <u>Males</u> |  |  |
| *ChAT-GAL4DBD, nsyb-QFAD, UAS-Luc* | 606 ± 56 | 5 |
| *tub-QS, ChAT-GAL4DBD, nsyb-QFAD, UAS-Luc* | 15 ± 1 | 5 |
| *ChAT-GAL4DBD, nsyb-QF2<sup>w</sup>AD, UAS-Luc* | 459 ± 6 | 5 |
| *tub-QS, ChAT-GAL4DBD, nsyb-QF2<sup>w</sup>AD, UAS-Luc* | 6.3 ± 0.3 | 5 |
| *ChAT-GAL4DBD, nsyb-p65AD, UAS-Luc* | 2803 ± 330 | 5 |
| *ChAT-GAL4DBD, nsyb-GAL4AD, UAS-Luc* | 27 ± 2 | 5 |
| *ChAT-GAL4DBD, UAS-Luc* | 5.8 ± 0.7 | 5 |
| *nsyb-QFAD, UAS-Luc* | 6.3 ± 0.7 | 5 |
| *nsyb-QF2<sup>w</sup>AD, UAS-Luc* | 8 ± 3 | 5 |
| *nsyb-GAL4QF, UAS-Luc* | 1377 ± 139 | 5 |
| *tub-QS, nsyb-GAL4QF, UAS-Luc* | 12 ± 1 | 5 |

**Supplementary Table 5.**
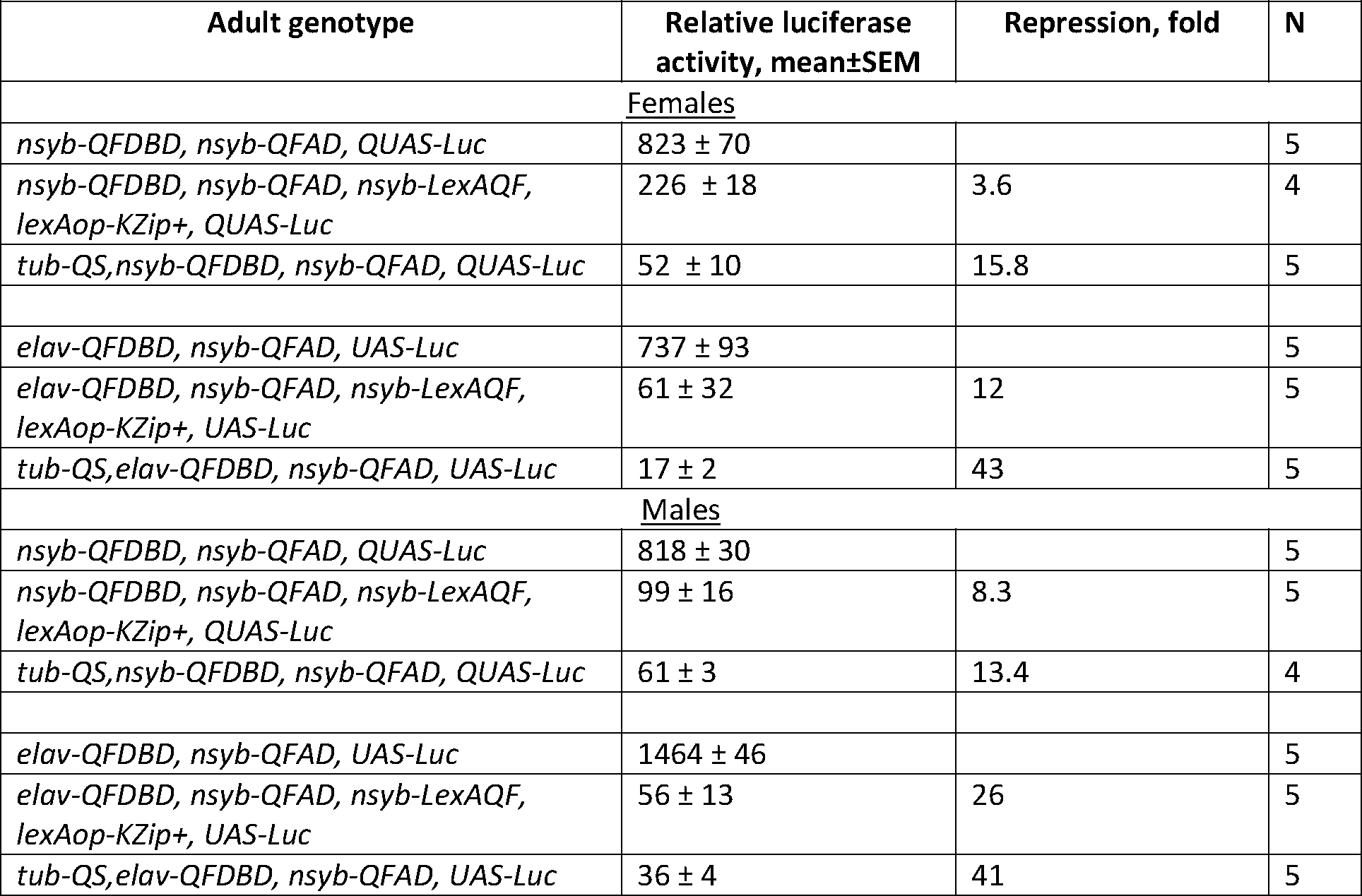
Quantification of repression by QS and KZip+

| Adult genotype | Relative luciferase activity, mean±SEM | Repression, fold | N |
| --- | --- | --- | --- |
| <u>Females</u> |  |  |  |
| *nsyb-QFDBD, nsyb-QFAD, QUAS-Luc* | 823 ± 70 | | 5 |
| *nsyb-QFDBD, nsyb-QFAD, nsyb-LexAQF, lexAop-KZip+, QUAS-Luc* | 226 ± 18 | 3.6 | 4 |
| *tub-QS, nsyb-QFDBD, nsyb-QFAD, QUAS-Luc* | 52 ± 10 | 15.8 | 5 |
| *elav-QFDBD, nsyb-QFAD, UAS-Luc* | 737 ± 93 | | 5 |
| *elav-QFDBD, nsyb-QFAD, nsyb-LexAQF, lexAop-KZip+, UAS-Luc* | 61 ± 32 | 12 | 5 |
| *tub-QS, elav-QFDBD, nsyb-QFAD, UAS-Luc* | 17 ± 2 | 43 | 5 |
| <u>Males</u> |  |  |  |
| *nsyb-QFDBD, nsyb-QFAD, QUAS-Luc* | 818 ± 30 | | 5 |
| *nsyb-QFDBD, nsyb-QFAD, nsyb-LexAQF, lexAop-KZip+, QUAS-Luc* | 99 ± 16 | 8.3 | 5 |
| *tub-QS, nsyb-QFDBD, nsyb-QFAD, QUAS-Luc* | 61 ± 3 | 13.4 | 4 |
| *elav-QFDBD, nsyb-QFAD, UAS-Luc* | 1464 ± 46 | | 5 |
| *elav-QFDBD, nsyb-QFAD, nsyb-LexAQF, lexAop-KZip+, UAS-Luc* | 56 ± 13 | 26 | 5 |
| *tub-QS, elav-QFDBD, nsyb-QFAD, UAS-Luc* | 36 ± 4 | 41 | 5 |

**Supplementary Table 6.**
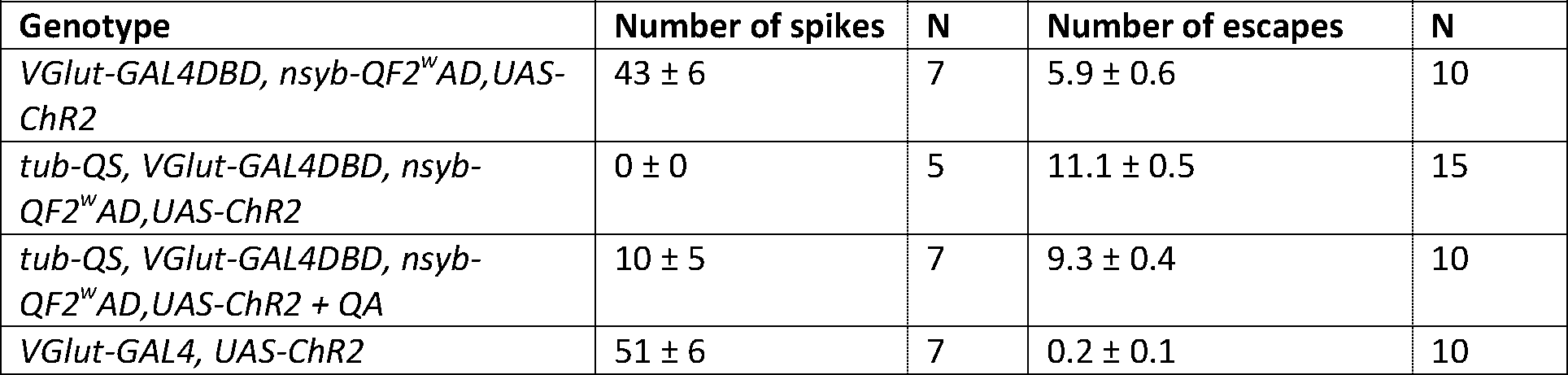
Optogenetic experiments in GAL4DBD + QF2^w^AD larvae

| Genotype | Number of spikes | N | Number of escapes | N |
| --- | --- | --- | --- | --- |
| *VGlut-GAL4DBD, nsyb-QF2<sup>w</sup>AD, UAS-ChR2* | 43 ± 6 | 7 | 5.9 ± 0.6 | 10 |
| *tub-QS, VGlut-GAL4DBD, nsyb-QF2<sup>w</sup>AD, UAS-ChR2* | 0 ± 0 | 5 | 11.1 ± 0.5 | 15 |
| *tub-QS, VGlut-GAL4DBD, nsyb-QF2<sup>w</sup>AD, UAS-ChR2 + QA* | 10 ± 5 | 7 | 9.3 ± 0.4 | 10 |
| *VGlut-GAL4, UAS-ChR2* | 51 ± 6 | 7 | 0.2 ± 0.1 | 10 |

**Supplementary Table 7.** Optogenetic experiments in split-QF larvae

| Genotype | Number of escapes | N |
| --- | --- | --- |
| *nsyb-QFDBD, nsyb-QFAD, QUAS-ChR2* | 0 ± 0 | 10 |
| *tub-QS, nsyb-QFDBD, nsyb-QFAD, QUAS-ChR2* | 7.4 ± 0.7 | 10 |
| *tub-QS, nsyb-QFDBD, nsyb-QFAD, QUAS-ChR2 + QA* | 0.3 ± 0.2 | 10 |
| *nsyb-QFDBD, nsyb-QF2<sup>w</sup>AD, QUAS-ChR2* | 0 ± 0 | 10 |
| *tub-QS, nsyb-QFDBD, nsyb-QF2<sup>w</sup>AD, QUAS-ChR2* | 8 ± 0.8 | 7 |
| *tub-QS, nsyb-QFDBD, nsyb-QF2<sup>w</sup>AD, QUAS-ChR2 + QA* | 0.1 ± 0.1 | 9 |

**Supplementary Table 8.** Thermogenetic experiments in split-QF adults

| Time,<br>min | Number of flies on the bottom of the vial |  |  |  |
| --- | --- | --- | --- | --- |
|  | *nsyb-QFDBD, nsyb-QFAD, QUAS-shibire<sup>TS</sup>* (N=5) | *tub-QS, nsyb-QFDBD, nsyb-QFAD, QUAS-shibire<sup>TS</sup>* (N=4) | *nsyb-QFDBD, nsyb-QF2<sup>w</sup>AD, QUAS-shibire<sup>TS</sup>* (N=5) | *tub-QS, nsyb-QFDBD, nsyb-QF2<sup>w</sup>AD, QUAS-shibire<sup>TS</sup>* (N=4) |
| 0 | 0 ± 0 | 0 ± 0 | 0.2 ± 0.2 | 0 ± 0 |
| 0.5 | 0.4 ± 0.2 | 0 ± 0 | 0 ± 0 | 0 ± 0 |
| 1 | 0.4 ± 0.2 | 0 ± 0 | 0 ± 0 | 0 ± 0 |
| 1.5 | 0.6 ± 0.4 | 0 ± 0 | 0 ± 0 | 0.5 ± 0.3 |
| 2 | 1.2 ± 0.6 | 0 ± 0 | 0.2 ± 0.2 | 0.25 ± 0.25 |
| 2.5 | 1.6 ± 0.8 | 0 ± 0 | 0.2 ± 0.2 | 0.25 ± 0.25 |
| 3 | 1.8 ± 0.8 | 0 ± 0 | 0.8 ± 0.6 | 0.25 ± 0.25 |
| 3.5 | 2.4 ± 1.1 | 0 ± 0 | 0.8 ± 0.6 | 0.5 ± 0.3 |
| 4 | 3.2 ± 1.2 | 0 ± 0 | 1 ± 0.5 | 0.75 ± 0.25 |
| 4.5 | 3.6 ± 1.3 | 0 ± 0 | 1.4 ± 0.7 | 0.75 ± 0.25 |
| 5 | 4 ± 1.3 | 0 ± 0 | 1.2 ± 0.7 | 0.75 ± 0.25 |
| 5.5 | 4.4 ± 1.4 | 0.25 ± 0.25 | 1.6 ± 0.7 | 0.5 ± 0.3 |
| 6 | 4.2 ± 1.4 | 0.25 ± 0.25 | 2 ± 0.7 | 0.75 ± 0.25 |
| 6.5 | 5.8 ± 1.4 | 0.25 ± 0.25 | 3 ± 1.2 | 1 ± 0.4 |
| 7 | 7 ± 1 | 0.25 ± 0.25 | 3 ± 1.4 | 0.5 ± 0.3 |
| 7.5 | 7.4 ± 0.8 | 0.25 ± 0.25 | 3.6 ± 1.3 | 0.5 ± 0.3 |
| 8 | 7.8 ± 0.7 | 0 ± 0 | 3.8 ± 1.3 | 0.5 ± 0.3 |
| 8.5 | 9 ± 0.5 | 0.25 ± 0.25 | 4.8 ± 1 | 0.5 ± 0.3 |
| 9 | 9.4 ± 0.2 | 0 ± 0 | 5.2 ± 1.3 | 0.75 ± 0.5 |
| 9.5 | 9.8 ± 0.2 | 0 ± 0 | 5.8 ± 1 | 0.75 ± 0.5 |
| 10 | 9.8 ± 0.2 | 0.5 ± 0.5 | 6.4 ± 1 | 1.3 ± 0.9 |
| 10.5 | 9.8 ± 0.2 | 0.5 ± 0.5 | 7.4 ± 1 | 1 ± 1 |
| 11 | 10 ± 0 | 0.25 ± 0.25 | 8 ± 0.6 | 0.5 ± 0.3 |
| 11.5 | 10 ± 0 | 0 ± 0 | 8.2 ± 0.7 | 0.25 ± 0.25 |
| 12 | 10 ± 0 | 0 ± 0 | 8.4 ± 0.7 | 0.3 ± 0.3 |
| 12.5 | 10 ± 0 | 0 ± 0 | 8.8 ± 0.5 | 0 ± 0 |
| 13 | 10 ± 0 | 0 ± 0 | 9.2 ± 0.5 | 0 ± 0 |
| 13.5 | 10 ± 0 | 0 ± 0 | 9.4 ± .4 | 0 ± 0 |
| 14 | 10 ± 0 | 0 ± 0 | 9.2 ± 0.5 | 0 ± 0 |
| 14.5 | 10 ± 0 | 0 ± 0 | 9.8 ± 0.2 | 0 ± 0 |
| 15 | 10 ± 0 | 0 ± 0 | 10 ± 0 | 0 ± 0 |

